# Timing of speech in brain and glottis and the feedback delay problem in motor control

**DOI:** 10.1101/2024.08.07.607082

**Authors:** John P. Veillette, Jacob Rosen, Daniel Margoliash, Howard C. Nusbaum

## Abstract

To learn complex motor skills, an organism must be able to assign sensory feedback events to the actions that caused them. This matching problem would be simple if motor neuron output led sensory feedback with a fixed, predictable lag. However, nonlinear dynamics in the brain and the body’s periphery can decouple the timing of critical events from that of the motor output which caused them. During human speech production, for example, phonation from the glottis (a sound source for speech) begins suddenly when subglottal pressure and laryngeal tension cross a sharp threshold (i.e. a bifurcation). Only if the brain can predict the timing of these discrete peripheral events resulting from motor output, then, would it be possible to match sensory feedback to movements based on temporal coherence. We show that event onsets in the human glottal waveform, as measured using electroglottography, are reflected in the human electroencephalogram during speech production, leading up to the time of the event itself. Conversely, glottal event times can be decoded from the electroencephalogram. After prolonged exposure to delayed auditory feedback, subjects recalibrate their behavioral threshold for detecting temporal auditory-motor mismatches, and decoded event times decouple from actual movements. This suggests decoding performance is driven by plastic predictions of peripheral timing, providing a missing component for hindsight credit assignment in motor control, in which specific feedback events are associated with the neural activity that gave rise to movements. We discuss parallel findings from the birdsong system suggesting that results may generalize across vocal learning species.

**Significance Statement:** To learn complex motor skills such as speech, the brain must pair actions with sensory feedback. However, feedback is delayed in time relative to actual movement, rendering this “hindsight credit assignment” problem a nontrivial task. We present evidence that events in the glottis, the articulatory organ that generates sound for human speech, are predictively encoded in the human brain during speech production. Given that corresponding events in auditory feedback are known to be prominently encoded in cortex, this highlights a common temporal marker by which articulatory gestures and sensory feedback could be aligned. Findings suggest activity in the human vocomotor system is shaped by biophysical dynamics in the vocal periphery, aligning with corresponding results in the birdsong system.

## Introduction

Speech production requires coordination of over a hundred muscles to achieve desired acoustic output (Simonyan and Horwitz, 2011). Were auditory feedback immediate, acoustic outcomes could be assigned to articulatory movements in real-time, allowing control policies to be learned in a “model-free” manner as in contemporary policy-based reinforcement learning systems (Kakade, 2001). However, muscle contraction, sensory transduction, and intervening multiple stages of neuronal transmission for both all result in intrinsic delays of the arrival of feedback to forebrain structures often associated with learned movements (Wolpert and Ghahramani, 2000; Wolpert et al., 2011). Consequently, vocal learning – and motor learning in general – is thought to require a “forward” model to actively predict current and future peripheral states and, in turn, the resulting sensory feedback from the recent motor output; discrepancies between predicted and actual feedback are used to update the forward model, which can then be leveraged offline to adjust an “inverse” model (i.e. control policy) which proposes movements to achieve desired states (Tourville and Guenther, 2011). This computational account has become a guiding framework for understanding both normal motor behavior and the symptomology of speech motor deficits such as stuttering (Bradshaw et al., 2021). But while forward model updates have been proposed to occur through reinforcement learning mechanisms (Sutton and Barto, 2018), the task of assigning outcomes to actions when feedback is temporally delayed – the hindsight credit assignment problem – remains a standing challenge in theoretical and applied reinforcement learning (Harutyunyan et al., 2019). In particular, when biomechanical dynamics at the body’s periphery are nonlinear, the critical events that generate sensory feedback may be temporally decoupled from the onsets of the muscle movements that caused them; in this case, motor neuron activity would not be separated from sensory feedback by a constant lag, and temporal coherence between the two would not be available as a cue for hindsight credit assignment.

Speech production is one such case. The physics of the vocal organ, the glottis, are such that acoustic output is highly discontinuous with respect to the physiological parameters – subglottal pressure and laryngeal tension – that are directly controlled by the brain (Titze, 1988; Sitt et al., 2008). In particular, in physical models of the glottis – or of the syrinx in songbirds, which has shared dynamics (Amador and Mindlin, 2014) – the position of the vocal folds oscillates given certain pressure/tension combinations but settles at a steady-state position otherwise; when pressure and tension crosses the boundary from a region that yields steady-state dynamics to one with oscillatory dynamics, the oscillations begin abruptly (see Fig. 1). These oscillations, called phonation or voicing, are what generate the pressure waves that pass through the upper vocal tract to be shaped into speech. Thus, while onsets of phonation and other discrete acoustic events at the glottis – referred to here as “glottal event onsets” – are directly caused by the motor output of the corticobulbar (laryngeal tension) and corticospinal (respiration/pressure) tracts, their correspondence to actual muscle movements is highly nonlinear.

**Figure 1:**
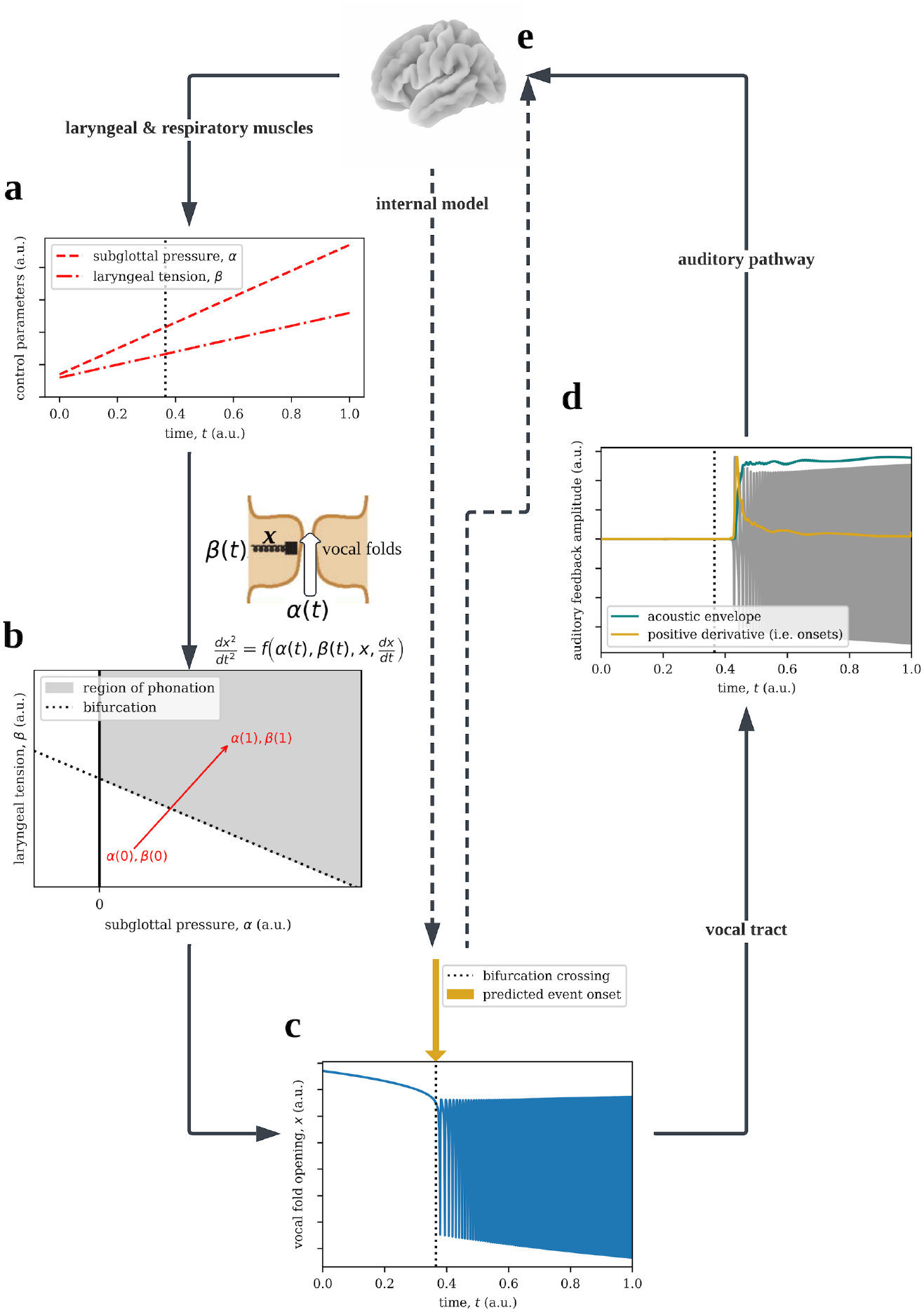
The necessity of prediction in the control of voicing. (**A**) Pictured are arbitrary, continuous trajectories of subglottal pressure and laryngeal tension, which the brain controls through the musculature. (**B**) In a single-mass model of the glottis, each vocal fold (a.k.a. “vocal chords”) are treated abstractly as a mass on a spring, where the stiffness of the spring is set by the tension and an opposing force by the pressure pushing up from the lungs, both interacting with the current position of the folds (e.g. pressure simply passes through the glottis when the folds are open). Per Newton’s law, the acceleration will be proportional to the sum of those position-dependent forces, and thus dynamics are governed by a second-order differential equation, which produces oscillatory behavior for some pressure/tension combinations but settles at a steady-state position otherwise (Titze, 1988). (**C**) When pressure/tension cross the bifurcating border between oscillation-producing and non-oscillatory combinations, phonation begins suddenly, here simulated from the pressure/tension trajectories in panel A using the normal form equations given by Mindlin (2013), which produces sound pressure waves that are filtered through the upper vocal tract to produce speech. To effectively control the onset of voicing, as to produce fluent speech, the brain must anticipate the time of the bifurcation crossing that triggers phonation. (**D**) The human auditory system is sensitive to the acoustic envelope, but also to the acoustic event onsets that directly result from bifurcation-crossings (Oganian and Chang, 2019), which we call glottal event onsets; however, this auditory feedback reaches the central nervous system at a non-negligible delay, even in normal speaking conditions. (**E**) If the brain has successfully anticipated the time of glottal events, feedback events will follow those predictions with a fixed lag, allowing sensory feedback to be accurately credited to articulatory events based on temporal coherence. Thus, we hypothesized that glottal event onsets – which we can observe non-invasively via electroglottograph – are reflected in cortex, separately from sensory feedback, during speech production even though they do not directly resemble neuromuscular output.

Some anticipatory mechanism, then, would be required for central neural activity to predict the timing of acoustic/phonatory events in the glottis. However, temporal coherence between such anticipatory representations and auditory feedback would greatly simplify the task of associating movements with their consequences. This proposition would potentially explain the longstanding but perplexing observation that playing a sine tone that is amplitude-modulated to match a human participants’ speech envelope with a delay has similarly deleterious effects on speech production as delayed auditory feedback (DAF) (Howell and Archer, 1984); such findings are not well explained by speech production models where forward prediction primarily pertains to the frequency content of sound (Guenther et al., 2006; Golfinopoulos et al., 2010). Consistent with this theory, auditory responses in human superior temporal gyrus prominently reflect acoustic event onsets (Oganian and Chang, 2019). However, cortical tracking of corresponding glottal event onsets – which can be measured noninvasively using electroglottography (EGG) – has yet to be demonstrated, which was the aim of the present study.

## Methods

### Methods Summary

We recorded subjects’ electroencephalography (EEG), audio, and glottis vibrations using electroglottograph (EGG) during speech production. We fit encoding models to test whether glottis event onsets explain EEG variance beyond previously identified features. Further, we exposed subjects to a period of DAF to, in principle, temporally shift their forward model predictions, reflected behaviorally in their threshold for detecting auditory feedback delays (Yamamoto and Kawabata, 2014). Finally, we tested whether a model trained to decode glottal events from brain activity prior to prolonged DAF exposure makes temporally shifted predictions after this threshold shift, reflecting subjects’ own updated predictions.

Throughout the experiment, subjects heard their auditory feedback through insert earphones while they spoke into a microphone. Subjects read sentences that are presented on a monitor in front of them during (1) a **baseline block** with no delay, i.e. normal speaking conditions, and (2) a **pretest block** in which we imposed a random delay between 0 and 250 ms in their auditory feedback on each trial. To assess whether glottal event times were encoded in the EEG, we trained a multivariate encoding model that predicted the continuous EEG from the speech audio envelope, speech audio onsets, the EGG envelope, and the EGG onsets, and we compared its predictive performance to a model that that excluded EGG audio onsets. Encoding models were trained during the pretest block, where audio and glottal events were temporally decoupled by design, and tested during the baseline block to assess generalization performance in normal speaking conditions.

Subjects then completed (3) an **exposure block** with a constant 200 ms delay, and (4) a **posttest block** that was identical to the pretest block. During the pretest and posttest blocks, subjects were asked to report, on each trial, whether they detected a delay in their auditory feedback; a multilevel logistic model was fit to find the threshold at which they could detect a delay 50% of the time in both blocks, so we could assess perceptual shift as a result of DAF exposure. Then, we trained a decoding model to predict glottal event onsets on the pretest block and tested it on the posttest block (and vice versa) to assess whether the predictions of models that achieve high decoding accuracy are systematically affected by exposure to delayed auditory feedback, consistent with decoding performance being driven by an internal representation that is plastic given sensorimotor experience.

### Ethics and Recruitment

We recruited 32 university students (ages 18-22; 15 males, 17 females) over our department’s study recruitment system. To obtain the desired level of precision for behavioral measurements, we used the following *a priori* criteria to determine sample size adaptively: subjects were recruited until the 90% HDPI of the Bayesian posterior for pre-to-post shift in delay detection threshold was less than 20 ms (regardless of whether the sign of the shift or whether the HDPI contained zero), which occurred after 29 subjects were collected, after which the additional 3 subjects who had already signed up for the experiment were run and data collection was concluded. Subjects self-reported being right handed, though 3 were considered ambidextrous by the Edinburgh Handedness Inventory (Oldfield, 2013). Subjects were excluded during recruitment if they reported having been previously diagnosed by a clinician with a hearing or speech deficit. Informed consent was obtained prior to the experiment, and procedures were approved by the Social and Behavioral Sciences Institutional Review Board at the University of Chicago (IRB21-1402).

### Experimental Design

After applying EEG and EGG electrodes to the subjects’ heads and necks, respectively, they sat in front of a monitor. In each trial of the subsequent task, subjects were asked to read one of the phonetically balanced Harvard Sentences, which was displayed on the monitor (Rothauser, 1969); sentences displayed were on average 9.1 (SD 1.2) syllables long and presented all at once. We controlled the trial-by-trial latency of auditory feedback, recorded through a microphone (Behringer ECM-8000) and played back through insert earphones (Etymotic ER-3C) which blocked out external sound, using a custom Python tool. Since such a system requires digitizing audio and then converting back to analog to play back to the subject, there is an intrinsic minimum hardware/software delay – that is, the actual delay between input and output audio when we specify a delay of zero – which we empirically measured as 4 milliseconds by recording input and output audio simultaneously on our EEG amplifier. For comparison, the intrinsic latency of other auditory feedback manipulation systems used in the literature tends to exceed 15 milliseconds (Kim et al., 2020). The delays recorded in our open dataset (and used in our analysis) have been corrected to account for this minimum delay (by simply adding 4 ms to the value of the “delay” field in the file where delays are recorded).

The task consisted of four blocks: (1) a baseline block in which subjects read 20 sentences with no imposed feedback delay, (2) a pretest block in which subjects read 50 sentences each with a random feedback delay, uniformly distributed between 0 and 0.25 seconds, (3) an exposure block in which subjects read 80 sentences with a constant 200 ms delay, and (4) a posttest block, identical to the pretest block. After each trial in pretest and posttest blocks, subjects were asked to indicate on a keyboard if they detected a delay between their movements and the sound of their voice. Sentence prompts were unique for each subject but were always phonetically balanced.

### Statistical Analysis

From both the delayed speech audio and the EGG, we computed spectrograms between 30 and 5000 Hz using a gammatone filter bank, which approximates the frequency response of the human cochlea and has been noted to outperform other sound frequency decompositions when predicting EEG from the speech envelope (Issa et al., 2024). The envelope of both the audio and EGG was computed by averaging the amplitude across equivalent rectangular bandwidths (i.e. an approximation to the frequency bandwidths in human hearing). As in prior EEG and electrocorticography work, an acoustic event onset time series was computed by taking the positive first derivative within each equivalent rectangular bandwidths and then averaging across them (Oganian and Chang, 2019; Brodbeck et al., 2023). Envelopes (but not onsets) were log-transformed, which better reflects the psychophysical curve for loudness and has been shown to improve prediction of EEG (Brodbeck et al., 2023). Subsequently, all features and the EEG were scaled to be of roughly equal variance using the *RobustScaler* in the *scikit-learn* package (Pedregosa et al., 2011). Feature scaling was performed within blocks, rather than across the whole dataset, to prevent train-test leakage during cross-validation.

For multivariate EEG encoding models, we estimated Temporal Response Functions as the coefficients of a linear ridge regression that predicts each electrode from values of the time-lagged audio and EGG envelopes from 0.5 seconds before to 0.5 seconds after each EEG time point using the *MNE-Python* package (Gramfort et al., 2014; Crosse et al., 2016). The regularization parameter for ridge regression was chosen using a grid search (between 0.1 and 1e7, log-spaced) to maximize leave-one-trial-out performance on the training set. While it has recently become popular in the EEG literature (Brodbeck et al., 2023), this approach has a longer history in single-cell physiology (Theunissen et al., 2001) and is also frequently used, though referred to as “voxelwise-encoding,” in the fMRI literature (Huth et al., 2016; Dupré la Tour et al., 2022). Encoding models were trained on the pretest block, where the randomly varied latency of the auditory feedback allowed the model to differentiate between the EGG and audio waveforms, which would otherwise be roughly synchronous, and then tested (i.e. cross-validated) under normal speaking conditions in the baseline block, where we computed the correlation between the predicted and actual values at each electrode to measure the encoding models’ generalization performance. We fit both an encoding model that includes all four audio and EGG features and one excluding EGG onsets to test whether EGG event onsets explain variation in brain activity beyond that already explained by the audio envelope, audio onsets, and the EGG envelope. Cross-validated correlations of both models’ predictions with each EEG electrode were statistically compared to chance using a permutation test where null permutations are generated by shuffling the predicted time-series across trials, and to each other using a paired permutation test, both corrected for multiple comparisons using threshold-free cluster enhancement (with clustering across neighboring electrodes) to control the familywise error rate (Smith and Nichols, 2009). Univariate (i.e. single feature) encoding models as in Fig. 2 were constructed by directly inverting our decoding models as described by Haufe and colleagues, such that they correspond exactly to the same patterns selected by our decoders (Haufe et al., 2014). We compare the electrode-by-time weights of the univariate encoding models to zero using the *t*-max procedure with 10,000 permutations to control for multiple comparisons, and we show the encoding model weights averaged across subjects in Fig. 2 (Nichols and Holmes, 2002). We also trained a separate encoding model for glottal event onsets on the posttest block, and we compared the weights of the pretest and posttest encoding models using a spatiotemporal cluster-based permutation test (Maris and Oostenveld, 2007).

**Fig. 2.**
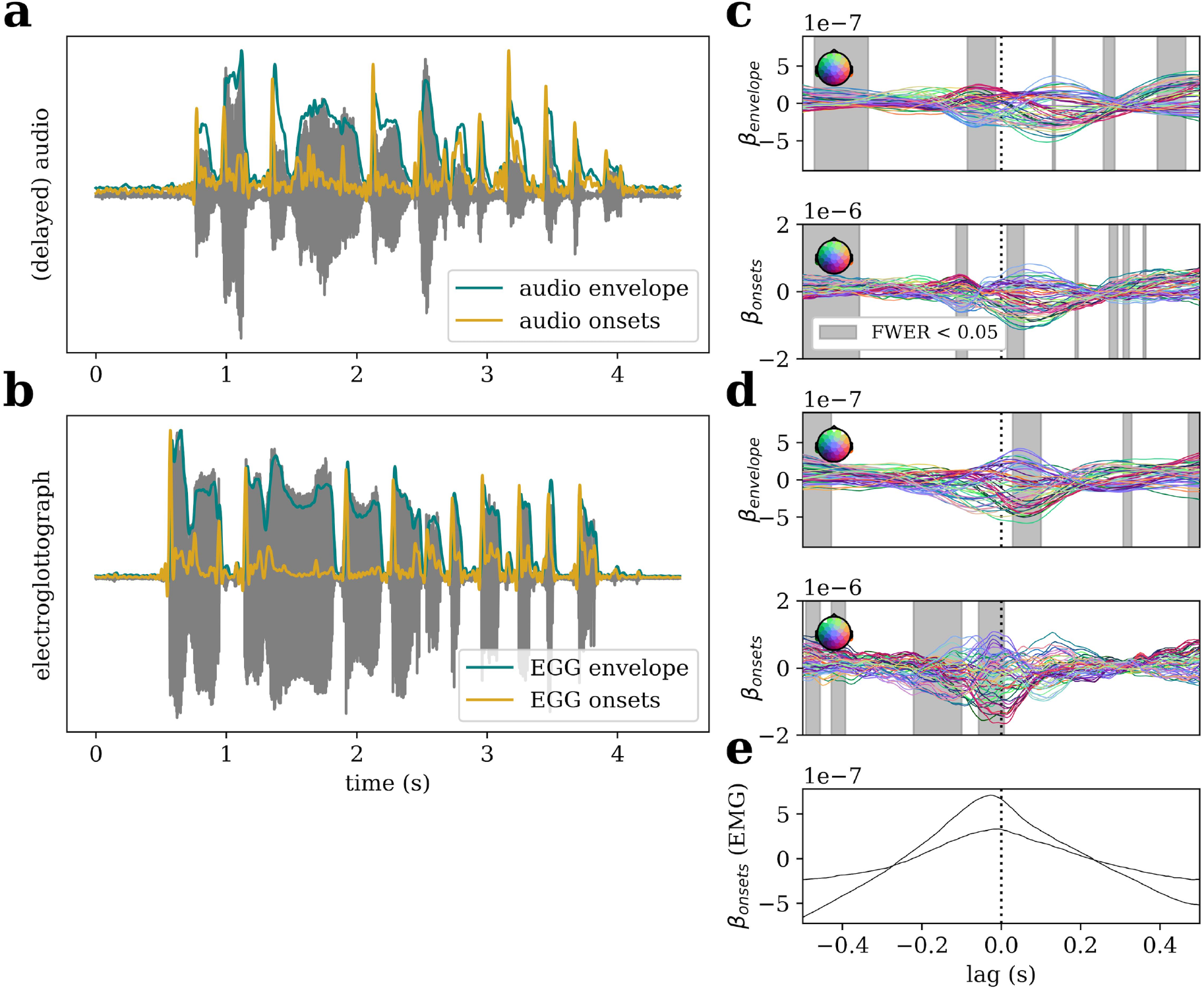
EEG temporal response functions to features of glottal waveform and auditory feedback waveform during speech production. From (**A**) the speech audio, played back to the subject at a delay subsequent to the baseline block, and from (**B**) the glottal waveform, we compute an amplitude envelope that approximates the frequency response of the cochlea and an “onset” time series that reflects the positive first derivative of the amplitude across all frequency bands – thus the onset of spectral events. Temporal response functions (TRFs), the EEG responses to these features of the (**C**) auditory feedback or (**D**) glottal waveform, are shown; a large coefficient in the TRF at, for example, lag 0.1 s means that a change in the value of that feature manifests in the EEG 0.1 seconds after that change. Shaded intervals cover time lags at which model weights for at least one electrode deviate from zero, after correcting for multiple comparisons. Model weights for encoding of glottal onsets are significant throughout an interval up to and including zero lag, consistent with neural activity anticipating glottal events, rather than merely responding to sensory feedback. (**E**) For comparison, a TRF to glottal onsets fit to the jaw EMG, rather than scalp EEG, channels. The predictions of the corresponding EMG-based decoding model are used as a control variable in subsequent EEG decoding analyses.

Decoding models were fit using the exact same procedure (same code) as the encoding models but in the reverse direction, predicting the values of each audio and EGG features from the -0.5 to 0.5 second time-lagged EEG electrodes. (That is, decoding models are like fitting an encoding model to the speech features, with the EEG electrodes as predictors.) While only the decoding of the EGG onsets is of interest in the present study, we report the cross-validated decoding performance – again, trained on pretest and tested on baseline – for all features, with *p*-values comparing each to chance using a permutation test where null permutations are generated by shuffling decoded time-series across trials.

Since EEG artifact correction methods can only be expected to attenuate, not completely remove, non-neural artifacts from the EEG signal, we performed a mediation analysis to assess statistically the extent to which EMG contamination may have explained successful decoding of glottal event onsets. To do this, we trained another decoding model to predict the glottal onsets directly from the two EMG channels on the pretest block, and generated predicted event onsets for the baseline block (exactly as we had with the EEG channels). For each subject, we then fit a mediation model for the linear relationship between the EEG-predicted onsets and the actual onsets, with the EMG-predicted onsets as a mediator; this gave us a per-subject “pure” estimate of the direct effect of the EEG in predicting onsets (left over when controlling for the confounding influence of EMG), and a separate estimate of the indirect effect which could be explained by EMG contamination. We compared the group-mean direct effect to chance with a permutation test (again shuffling the EEG-predicted time series across trials). We also performed a permutation test separately for each subject and fit a *p*-curve mixture model to the resultant *p-*values, which allowed us to use model comparison to assess relative evidence for population prevalence models in which there is a nonzero direct effect in none of, all of, or merely some subjects in the population (Veillette and Nusbaum, 2025); specifically, Bayesian model comparison is performed using the expected log pointwise predictive density (ELPD) with Pareto-smoothed importance sampling leave-one-out-cross-validation (Vehtari et al., 2017). We report bootstrap 95% confidence intervals for the proportion of the total effect that is accounted for by the direct effect.

To manipulate the content of subjects’ forward models, we exposed subjects to 0.2 second delayed auditory feedback throughout the exposure block, after which the same procedure as the pretest block was repeated in posttest. We estimated the shift in subjects’ threshold for detecting auditory-motor asynchronies, where “threshold” is defined as where the probability of detection is 50%, using a Bayesian multilevel logistic regression estimated with the *PyMC* package (Patil et al., 2010). Under the assumption that subjects detect a delay when their auditory feedback is sufficiently different from the feedback they predict, this threshold is a behavioral index of the content of their forward model and, as a corollary, a pretest to posttest change in threshold indicates a change in their forward model. Since delaying auditory feedback impairs speech production, usually resulting in slowing of speech or other disfluencies, we also fit a multilevel Poisson regression for speech rate as a function of delay and any change in that function from the pretest to posttest block. The speech rate on each trial was measured as the number of syllables in the target sentence divided by the duration of voicing, which was quantified in *Praat*.

If the EEG signal that reflects EGG event onsets is indeed an index of predictive cortical tracking of glottal events then one might expect that signal should change from pretest to posttest as do subjects’ detection thresholds. To test this hypothesis, we take those models which have been trained to predict EGG event onsets in the pretest block, and we generate its predicted onset time series for the posttest block; we then compute the time lag at which the cross-correlation between the predicted and actual event onsets is maximal as an estimate of how much predictions have shifted in time between pretest and posttest. Additionally, we train a decoding model on the posttest block and compute the same cross-correlation lag testing on the pretest block.

These measurements allow us to assess the possibility that the signal driving decoding is (a) a response to the glottal waveform via another sensory modality, e.g. somatosensory, or through bone conducted auditory feedback, or it reflects (b) a movement/muscle artifact or contamination from the EGG recording itself. Should either of these possibilities account for above-chance decoding performance, then there should be no trend in the central tendency of pre-to-posttest prediction lags or post-to-pretest lags with increasing decoder performance; at-chance models will always average zero lag, since any peak in the cross-correlation function would be spurious, and higher performing models will tend toward zero lag as signal-to-noise for decoding from the confound signal, which should not be affected by perceptual recalibration following DAF, improves. Conversely, if the signal that drives decoding performance is related to subjects’ forward model, then the predictions of decoders with higher signal-to-noise should be more affected (since low performing models will still be randomly distributed about zero lag, as their predictions are essentially random). Thus, the alternative hypothesis predicts a monotonic trend in pre-to-posttest lags and an opposite trend in post-to-pretest lags as baseline decoding performance improves. Thus, we test the null hypothesis using a Spearman rank correlation between the cross-validated correlation (trained on pretest, tested on baseline) for each subject and their pre-to-posttest shift in decoded onsets. We additionally report rank correlations between the lags and the direct (EEG) and indirect (EEG) effect estimates.

It is not necessarily the case that a shift in detection threshold corresponds directly to a same-direction shift in the expected time of glottal events as predicted by subjects’ forward models; psychophysical evidence indicates that humans also maintain a prior on the latency between action and sensory outcomes, and whether the expected movement time or expected action-outcome latency is updated following a prediction error would depend on the relative prior precision of the two estimates (Legaspi and Toyoizumi, 2019). However, the former explanation does make the distinct prediction that that pretest-trained decoder predictions will be delayed in the posttest block, and posttest-trained decoder predictions will be early in the pretest block. To test this additional prediction, we aimed to quantify the average temporal shift in decoded glottal event onsets from before to after adaption while controlling for the confounding influence of EMG contamination, which is indexed by the indirect effect estimate from the mediation analysis described above. To do so, we fit a robust linear regression of the form l*a*g = α_0_ + α_1_ (*indi*r*e*c*t e*ff*e*c*t*), such that the intercept α_0_ is interpreted as the expected value of the lag when the indirect effect is equal to zero. We report the regression *p-*value and confidence interval for this intercept term, fit to (a) just the pre-to-post lags and (b) the average lag for each subject, where post-to-pre lags have been flipped so their sign has the same interpretation as the pre-to-post lags.

### EEG Acquisition and Preprocessing

EEG was acquired at 10,000 Hz using an actiCHamp amplifier (Brain Products, GmbH) with 60 active EEG electrodes, two electrooculogram (EOG) electrodes below the eyes, and two electromyogram (EMG) channels over the jaw muscles in front of the ears, referenced to their ipsilateral mastoid. The precise positions of each EEG electrode were measured and recorded using a CapTrak (Brain Products, GmbH).

Since subjects were speaking during the study, much care was taken to attenuate contamination of the EEG signal from facial muscles. EEG was first processed using the PREP pipeline, which performs bad channel identification/interpolation, re-referencing, and line-noise removal in a standardized, robust manner (Bigdely-Shamlo et al., 2015; Appelhoff et al., 2022). Subsequently, we bandpass-filtered the EEG between 1-70 Hz and then decomposed the data into 60 independent components (ICs) and removed components that were significantly correlated with the EOG channels. We also, importantly, removed ICs that showed evidence of facial EMG contamination using the spectral slope, peripheral (i.e. far from vertex) power, and focal point (i.e. low spatial smoothness) criteria previously validated by Dharmaprani and colleagues using data from paralyzed patients (Dharmaprani et al., 2016). After removal of artifactual ICs, we then epoched the data and dropped trials that still showed evidence of strong non-neural signal contamination (i.e. peak-to-peak voltage exceeding 150 microvolts). To address any subtler movement artifacts, we then computed the current source density transformation of the EEG data using a spherical spline surface Laplacian. This transformation, effectively a spatial high pass filter, attenuates the effects of volume conduction across the scalp such that each electrode predominantly reflect nearby physiological sources (Kayser and Tenke, 2015). Previous work, again using data from paralyzed patients, has shown that the surface Laplacian effectively removes EMG contamination from central and peri-central electrodes (Fitzgibbon et al., 2013). Finally, the EEG was down-sampled to 140 Hz (i.e. twice the lowpass cutoff) to reduce the computational burden of our analysis.

All EEG preprocessing was automated to ensure reproducibility – using *pyprep*’s implementation of the PREP pipeline and the *MNE-Python* implementation of ICA and routines to select artificial ICA components –but data quality was manually verified after the PREP pipeline and again after artifact correction. In addition to the preprocessed data itself, detailed preprocessing reports that were generated at these points are available for each individual subject as part of our open dataset. These reports contain power spectra of the EEG signal after PREP, a list of electrodes that were interpolated (and the criteria by which they were flagged during PREP), visualized scalp topographies of all removed ICs, and remaining trial counts for each condition after rejection criteria are applied.

### EGG and Audio Acquisition and Synchronization

EGG was acquired using an EG2-PCX2 electroglottograph amplifier (Glottal Enterprises, Inc.) and audio using a Behringer ECM-8000 microphone with a roughly flat frequency response. These signals were sent from the EGG amplifier to a StimTrak (Brain Products, GmbH) via an audio passthrough cable, and were then digitized by our EEG amplifier so that all signals were recorded on the same clock. Like most EGG systems, the EG2-PCX2 has a built-in highpass filter, which introduces phase-specific delays into the recorded signal. To reverse this phase delay, we applied a digital equivalent of this 20 Hz highpass filter backwards after acquisition. (This sequential forward-backward filtering is the same operation that “zero phase shift” filters, like *scipy*’s and MATLAB’s *filtfilt* functions, apply.)

### Data and Code Availability

All raw data and preprocessed derivatives, included fitted models, are available on *OpenNeuro* (https://doi.org/10.18112/openneuro.ds005403.v1.0.1) and are organized according to the Brain Imaging Data Structure conventions for EEG (Pernet et al., 2019). Versioned analysis code is archived on *Zenodo* (https://zenodo.org/doi/10.5281/zenodo.13238912), and task code is available on GitHub (https://github.com/apex-lab/daf-experiment/releases/tag/v1.0.0).

## Results

### Glottal waveform events reflected in the EEG during speech production

The encoding model that incorporates all features – speech audio envelope, audio onsets, EGG envelope, and EGG onsets – performed robustly above chance (TFCE statistic = 11.34, p = 0.0001), as did the model that excludes EGG onsets (TFCE = 10.12, p = 0.0001). However, including the EGG onsets in the model did improve out-of-sample prediction of the EEG signal (TFCE = 2.38, p = 0.0085), indicating that the time of events in the EGG explains additional variance in cortical neural activity that is not explained by these other features.

The EEG temporal response function (TRF) to EGG onsets, visualized in Fig. 2, shows significant model weights leading up to and including the actual time of the glottal event. This is too early to be caused by sensory feedback (which, by definition, occurs after the event) and aligns with the theoretically predicted timescale of a forward model’s predictive representations, evidence of which is observed both in the motivating songbird studies and, though outside of the auditory system, in primates (Mulliken et al., 2008; Amador et al., 2013; Dima et al., 2018). However, a TRF time-locked to movement could also be predicted by either a movement artifact or perhaps by a motor command precipitating the movement itself; our subsequent decoding analyses were designed to assess the possibility of such confounds.

### Glottal event onsets decoded from the EEG

All four features, the envelope and onsets for audio and EGG signals, could be decoded from the EEG with above-chance cross-validation performance, with all *p* = 0.0001, the minimum possible *p*-value for our permutation test. The out-of-sample correlation [95% CI] between decoded and actual feature time series was *r* = 0.233 [0.199, 0.268] for the audio envelope, *r* = 0.101 [0.086, 0.117] for audio onsets, *r* = 0.349 [0.312, 0.382] for EGG envelope, and *r* = 0.136 [0.120, 0.152] for EGG onsets. The full distribution of subject-level decoding performances is shown in Fig. 3a, next to the encoding model performances in Fig. 3b-c. Moreover, when we control for EMG contamination in a mediation analysis, we estimate that 74.5% [65.8, 80.8] of the total linear relationship between EEG-predicted glottal onsets and actual onsets is accounted for by a direct effect (i.e. not by EMG), and our permutation test found this direct effect was significant at the group level (*p* = 0.0001). We also performed a permutation test for each subject individually to estimate the population prevalence of the effect with a *p*-curve mixture model (Veillette and Nusbaum, 2025); Bayesian model comparison found that data were best explained by a model in which all subjects (rather than just some) showed a direct effect (ELPD = 153.2), assigning a weight of 0.95 to that model. (This is interpreted loosely as a 0.95 posterior probability that this model would best predict the results of an additional subject.) In other words, formal prevalence inference suggests this effect may be nearly universal in the population from which we sampled (Veillette and Nusbaum, 2025).

**Fig. 3.**
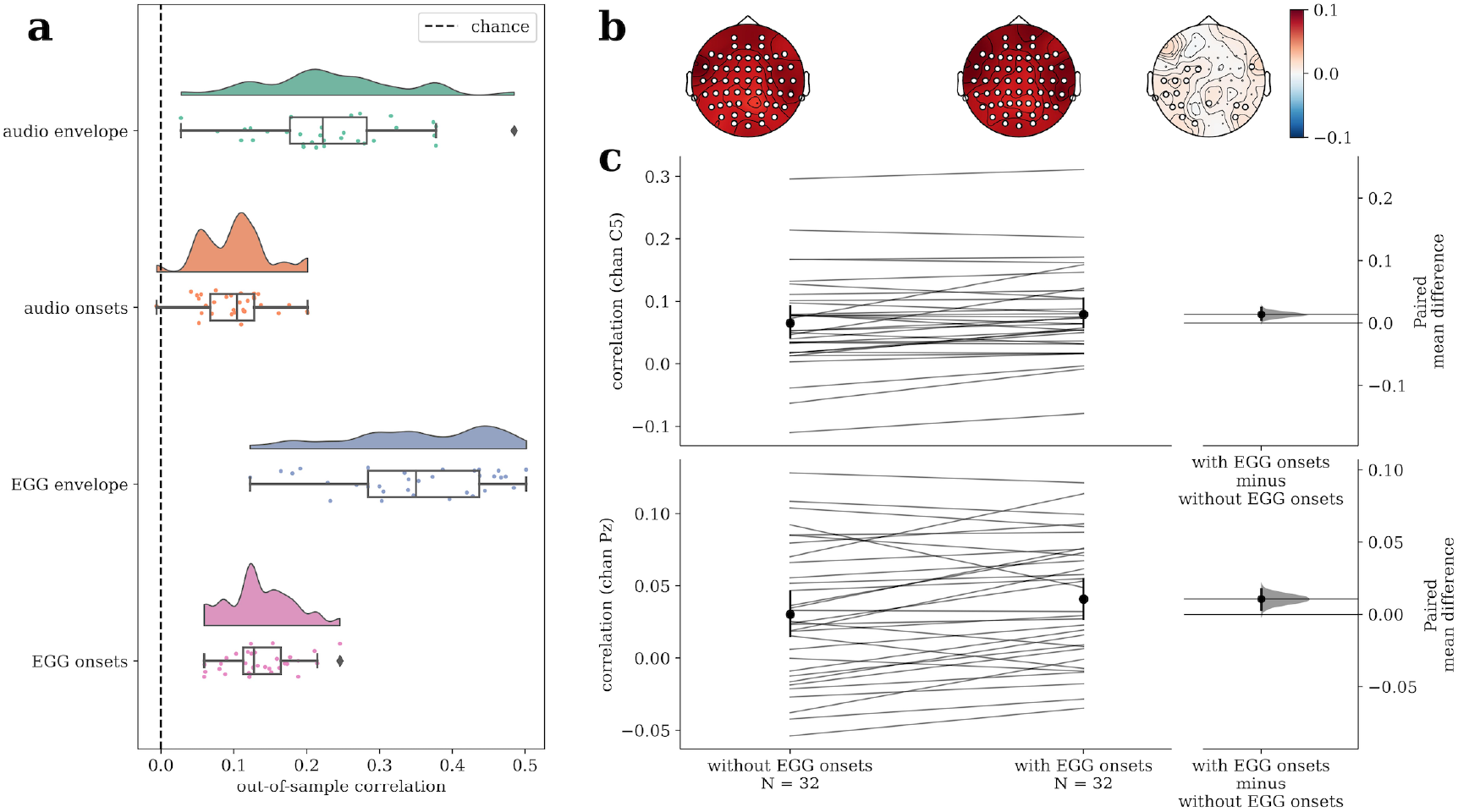
Out-of-sample predictive performance of decoding and encoding models. (**A**) Per-subject cross-validated Pearson correlations between actual time series of the audio/EGG features and those decoded from the EEG are shown as individual points, with overlaid boxplots showing quartiles of the observed correlations and whiskers extend 150% the interquartile range past the upper and lower quartiles to show the realistic extent of the distribution. Diamonds represent subjects falling outside of this range, and kernel density estimates are shown above. (**B**) The cross-validated Pearson correlation between actual EEG and that predicted from the audio/EGG features, shown for all electrodes. The model on the left excludes the EGG onsets as a predictor, the middle includes EGG onsets, and the difference in performance is shown on the right. White circles mark electrodes that are significant after multiple comparisons correction. (**C**) The same encoding model performances are shown for a central (C5) and a parietal (Pz) electrode; paired correlations for each subject, their bootstrap means with 95% confidence intervals, and the bootstrap distribution of the mean differences are shown. All decoding and encoding models visualized here were trained in the pretest block (i.e. before exposure) and tested in the baseline block – in other words, in normal speaking conditions.

### Temporal recalibration of perception of auditory-motor synchrony after prolonged exposure to delayed auditory feedback

The group-level parameters [95% highest posterior density interval (HPDI)] of the logistic multilevel model (time units in milliseconds) used to measure delay detection threshold were β _*intercept*_= -2.19 [-2.64, -1.70], β_*delay*_ = 36.6 [31.16, 42.25], β_*adaption*_ = -0.97 [-1.52, -0.414], β_*delay*×*adaption*_ = 2.61 [-3.18, 8.47]. The detection threshold, or the delay at which the probability of detection is 50%, shifted from mean 59.8 ms pre-exposure to 80.8 ms post-exposure; in other words, subjects’ detection thresholds shifted by about 20.9 ms [10.7, 31.6] after the exposure block. Trial-level behavioral data (Fig. 4, top), group-level logistic curves, and both group- and subject-level estimates of threshold shift (Fig. 4, bottom) are visualized in Fig. 4. Recalibration of auditory-motor asynchrony detection thresholds indicates that subjects’ internal predictions about the time of articulatory events and/or the latency of their sensory consequences were altered by the DAF manipulation.

**Fig. 4.**
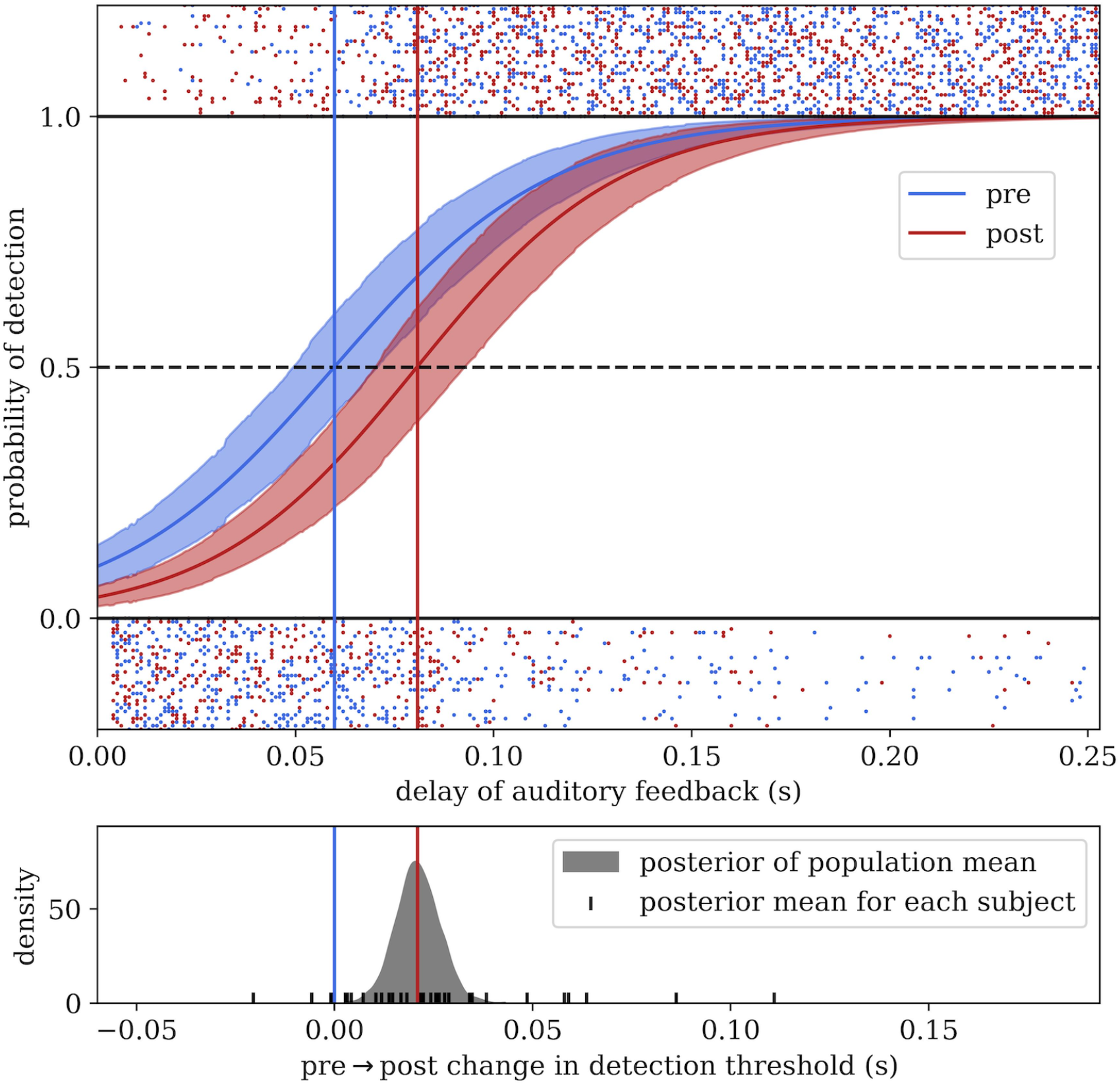
Behavioral shift in delay detection threshold following exposure to delayed auditory feedback. (top) The group-level logistic regression functions describing the probability of detecting an auditory-motor asynchrony in the pretest and posttest blocks, which flank an exposure block where subjects are exposed to a constant auditory feedback delay. Trial-by-trial responses are shown above and below the logistic curves, with rows corresponding to subjects. (bottom) The posterior distribution of the population/group average change in the delay detection threshold, with point estimates shown for each individual subject.

In contrast, there was not strong evidence of motor adaption as a consequence of prolonged DAF exposure. A Poisson regression found that speech rates indeed slowed, as expected, as a function of the trial-by-trial delay (β_*inte*rc*ept*_= 1.40 [1.35, 1.45], β_*delay*_ = -0.87 [-1.15, -0.61]). However, the credible intervals of adaption-related changes both contained zero (β_*adaption*_ = 0.04 [-0.02, 0.010], β_*delay*×*adaption*_ = -0.03 [-0.38, 0.29]). Taking the upper edge of the 95% credible interval as an upper bound, this suggests any main effect of exposure is smaller than a 0.37 syllables/second difference, if such an effect exists at all.

### Decoded glottal event times affected by perceptual recalibration

When we trained decoding models to predict EGG onsets from brain activity on the pretest block, we found that models which performed better in normal speaking conditions (i.e. baseline block) predicted later onset times relative to the actual EGG events when tested during posttest (Spearman’s rank correlation = 0.410, p = 0.0198). The temporal shift was also positively correlation with the EEG direct effect (rank correlation = 0.396, p = 0.0251) but negatively correlated with EMG contamination (rank correlation = -0.446, p = 0.0105). In contrast, models trained on models trained on posttest (which resulted in different model weights; cluster based permutation test: p *=* 0.0039, see Fig. 5b) and tested at pretest showed the exactly the opposite pattern, where lags varied negatively with model score (rank correlation = -0.350, p = 0.0497) and with the direct EEG effect (rank correlation = -0.413, p = 0.0189), though not significantly with the indirect effect (rank correlation = 0.192, p = 0.293). The fact that decoder performance is monotonically related to how much a decoder is affected by exposure to an auditory-motor delay rules out the possibility that decoding performance is explained by a movement artifact or non-plastic sensory response, which would be invariant to temporal recalibration. (We do note that, despite this group trend, lags of zero are still common in the sample, so it is certainly possible that such confounds still contribute to decoding performance as also suggested by our mediation analysis – they just cannot fully explain it.)

**Figure 5:**
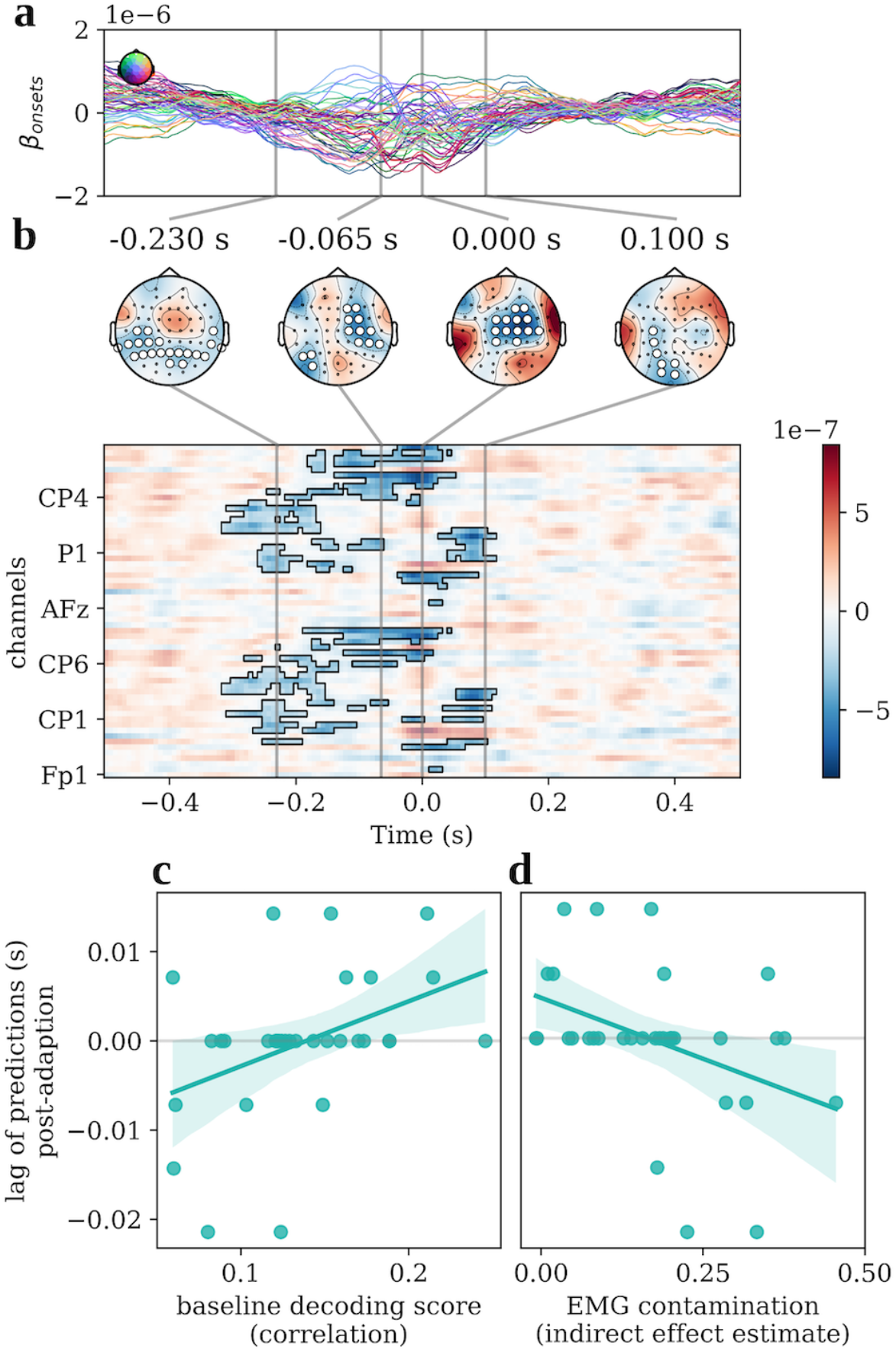
Exposure to delayed auditory feedback alters the neural activity tracking glottal event times. (**A**) The weights of EEG encoding model for glottal event onsets trained during the posttest block. (**B**) Difference wave for posttest encoding model vs. the encoding model trained during pretest and seen in in Fig. 2d, with spatiotemporal cluster that exceeded change magnitude in cluster-based permutation test highlighted and electrodes marked on scalp. (**C**) The lag between actual event times and event times predicted by decoder trained in the pretest block and tested in the posttest block, shown as a function of baseline decoding performance and (**D)** as a function of estimated EMG contamination. High performing models and models that show minimal EMG contamination tend toward positive shifts, suggesting EEG-based predictions are delayed relative to events following prolonged exposure to delayed auditory feedback.

To estimate the mean temporal lag in predictions due to DAF exposure, we performed a robust regression while controlling for the confounding influence of EMG contamination (see Fig. 5d). Using only the pre-to-posttest lags, we estimated that shift as roughly 3.6 milliseconds (p = 0.0359, 95% CI: [0.2, 7.0]). Incorporating both pre-to-posttest and (sign-flipped) post-to-pretests lags as repeated measures, we estimated a slightly smaller but still significant shift of roughly 2.9 milliseconds (p = 0.0483, 95% CI: [0.02, 5.8]). In other words, we do find evidence of a slight positive shift on average, consistent with an internal prediction of event time updating after exposure to delayed sensory feedback. However, we note that the estimated shift is an order of magnitude smaller than the behavioral shifts in the delay-detection threshold, and thus unlikely to fully explain perceptual adaption.

As the decoded signal can decouple in time from movements, it is neither well explained by a motor command as traditionally conceptualized, which would lead the movement with a constant delay, nor by a movement artifact, which would be inherently synchronous to the movement. Rather, high decoding performance appears to be driven by a signal that is ordinarily temporally coherent with glottal articulatory movements but can decouple following exposure to an auditory-motor mismatch. These operating characteristics are indicative of an internal prediction, which is malleable to sensorimotor experience.

## Discussion

The present study presents evidence that neural activity in human cortex tracks the timing of events in the glottal waveform during speech production in roughly temporally coordinate fashion. Acoustic events in the glottal waveform have a lawful temporal correspondence to the subglottal pressure and laryngeal tension gestures that are used to control the vibration of the glottis (Titze, 1988), as is true for the equivalent structures in avian vocal learners (Perl et al., 2011; Mindlin, 2013; Boari et al., 2015). Moreover, the glottal waveform serves as the acoustic source that, once filtered through the upper vocal tract, results in the speech waveform (Titze, 1998). Thus, under normal speaking conditions, event onsets in the glottal waveform have a one-to-one correspondence with subsequent acoustic events in the auditory stream, and such onset events are known to be prominently reflected in the auditory cortical representation of speech (Oganian and Chang, 2019). In other words, cortical tracking of glottal gestural/acoustic event times could contribute to the circuit resolving the temporal credit assignment problem – based on temporal coherence between event times – in the control of voicing (Harutyunyan et al., 2019).

Work in birdsong notably mirrors but also helps to elaborate these observations. In the HVC (premotor/association cortex analog), certain neurons produce spike bursts and interneurons are suppressed within milliseconds of transitions between the syringeal pressure and tension gestures used to control birds’ analog to the human glottis within their syrinx (Amador et al., 2013). As bursts occur too late to have triggered movements but too early to represent sensory feedback, this synchronous activity has been interpreted as a state prediction. More recent work has shown that some primary motor cortex analog projecting HVC neurons also burst at syllable onsets or offsets and at prominent transitions within syllables, although the timing of these burst relative to syringeal gestures has yet to be formally evaluated (Moll et al., 2023). Some models of birdsong production proposed that this sort anticipated synchronization of motor neuron activity with peripheral events emerges organically as a characteristic of certain coupled dynamical systems, which yield zero lag relationships between hierarchically organized structures (Alonso et al., 2015; Dima et al., 2018). The patterns of respiratory activity in singing canaries are also predicted by such models (Alonso et al., 2015; Dima et al., 2018). Notably, pressure-tension gesture transitions are directly reflected in the derivative of the song envelope (Boari et al., 2015), and some Field L (auditory cortex analog) neurons selectively respond to the same derivative information in auditory stimuli (Sharpee et al., 2011). As some HVC neurons are prone to repeat bursts after fixed intervals following their initial recruitment during song, rebound bursts may occur near the time of sensory feedback (Daou and Margoliash, 2020; Fetterman and Margoliash, 2023). These observations provide elements for simultaneous neural representation of gestures and their acoustic outcomes, which could be linked by well-documented coincidence detection mechanisms (Joris et al., 1998; Matell and Meck, 2004).

It is worth elaborating that such a mechanism, emphasizing event onsets and transitions rather than event content as described, complement the content-specific forward model predictions posited by extant models of motor control (Tourville and Guenther, 2011). Frequency-specific suppression of sounds resulting from movement is well established in mammalian auditory cortex (Schneider and Mooney, 2018; Schneider et al., 2018), but to learn such content-specific mappings in the first place, we posit an organism must grapple with the hindsight credit assignment problem. Representing articulatory movements in a manner that is temporally coherent with resultant peripheral events ensures sensory feedback occurs at a fixed lag relative to motor neuron activity, which affords learning content-specific action-outcome mappings via simple associative mechanisms. In speech production, this would predict – as we present evidence for here – a temporal correspondence between some neural activity and the onsets of discrete acoustic events that occur when continuous changes in subglottal pressure and laryngeal tension act on the nonlinear physical dynamics of the voice’s sound source, the glottis (Sitt et al., 2008; Boari et al., 2015). In this view, then, hindsight credit assignment when processing auditory feedback is emergently facilitated by the manner in which the central nervous system coordinates with the vocal organ, circumventing the need for specialized CNS mechanisms to temporally realign actions-outcome pairs post hoc.

The authors declare no conflict of interest.

## Author Contributions

Conceptualization: JPV, DM, and HCN; Data curation: JPV and JR; Formal analysis: JPV; Funding acquisition: DM and HCN; Investigation: JPV and JR; Methodology: JPV; Project administration: JPV; Resources: DM and HCN; Software: JPV; Supervision: HCN; Validation: JPV; Visualization: JPV; Writing - original draft: JPV; Writing - review & editing: JPV, JR, DM, and HCN.

## Acknowledgements

National Science Foundation grant BCS 1835181 (DM and HCN); National Science Foundation fellowship DGE 1746045 (JPV); Neubauer Family Distinguished Doctoral Fellowship (JPV).

## Notes

https://doi.org/10.18112/openneuro.ds005403.v1.0.1

https://zenodo.org/doi/10.5281/zenodo.13238912

### Competing Interest Statement

The authors have declared no competing interest.

### Summary of Updates

Updated with revisions made during peer review.

## References

Alonso RG, Trevisan MA, Amador A, Goller F, Mindlin GB (2015) A circular model for song motor control in Serinus canaria. Front Comput Neurosci 9 Available at: https://www.frontiersin.org/journals/computational-neuroscience/articles/10.3389/fncom.2015.00041/full [Accessed August 6, 2024].

Amador A, Mindlin GB (2014) Low dimensional dynamics in birdsong production. Eur Phys J B 87:1–8.

Amador A, Perl YS, Mindlin GB, Margoliash D (2013) Elemental gesture dynamics are encoded by song premotor cortical neurons. Nature 495:59–64.

Appelhoff S, Hurst A, Lawrence A, Li A, Mantilla R, Yorguin J, O’Reilly C, Xiang L, Dancker J (2022) PyPREP: A Python implementation of the preprocessing pipeline (PREP) for EEG data. Available at: 10.5281/zenodo.6363575 [Accessed February 16, 2023].

Bigdely-Shamlo N, Mullen T, Kothe C, Su K-M, Robbins KA (2015) The PREP pipeline: standardized preprocessing for large-scale EEG analysis. Front Neuroinformatics 9 Available at: https://www.frontiersin.org/articles/10.3389/fninf.2015.00016 [Accessed February 16, 2023].

Boari S, Perl YS, Amador A, Margoliash D, Mindlin GB (2015) Automatic reconstruction of physiological gestures used in a model of birdsong production. J Neurophysiol 114:2912–2922.

Bradshaw AR, Lametti DR, McGettigan C (2021) The Role of Sensory Feedback in Developmental Stuttering: A Review. Neurobiol Lang 2:308–334.

Brodbeck C, Das P, Gillis M, Kulasingham JP, Bhattasali S, Gaston P, Resnik P, Simon JZ (2023) Eelbrain, a Python toolkit for time-continuous analysis with temporal response functions Martin AE, Shinn-Cunningham BG, Slaats S, eds. eLife 12:e85012.

Crosse MJ, Di Liberto GM, Bednar A, Lalor EC (2016) The Multivariate Temporal Response Function (mTRF) Toolbox: A MATLAB Toolbox for Relating Neural Signals to Continuous Stimuli. Front Hum Neurosci 10 Available at: https://www.frontiersin.org/journals/human-neuroscience/articles/10.3389/fnhum.2016.00604/full [Accessed July 15, 2024].

Daou A, Margoliash D (2020) Intrinsic neuronal properties represent song and error in zebra finch vocal learning. Nat Commun 11:952.

Dharmaprani D, Nguyen HK, Lewis TW, DeLosAngeles D, Willoughby JO, Pope KJ (2016) A comparison of independent component analysis algorithms and measures to discriminate between EEG and artifact components. In: 2016 38th Annual International Conference of the IEEE Engineering in Medicine and Biology Society (EMBC), pp 825–828. IEEE.

Dima GC, Copelli M, Mindlin GB (2018) Anticipated Synchronization and Zero-Lag Phases in Population Neural Models. Int J Bifurc Chaos 28:1830025.

Dupré la Tour T, Eickenberg M, Nunez-Elizalde AO, Gallant JL (2022) Feature-space selection with banded ridge regression. NeuroImage 264:119728.

Fetterman GC, Margoliash D (2023) Rhythmically bursting songbird vocomotor neurons are organized into multiple sequences, suggesting a network/intrinsic properties model encoding song and error, not time. :2023.01.23.525213 Available at: https://www.biorxiv.org/content/10.1101/2023.01.23.525213v1 [Accessed July 15, 2024].

Fitzgibbon SP, Lewis TW, Powers DMW, Whitham EW, Willoughby JO, Pope KJ (2013) Surface Laplacian of Central Scalp Electrical Signals is Insensitive to Muscle Contamination. IEEE Trans Biomed Eng 60:4–9.

Golfinopoulos E, Tourville JA, Guenther FH (2010) The integration of large-scale neural network modeling and functional brain imaging in speech motor control. NeuroImage 52:862–874.

Gramfort A, Luessi M, Larson E, Engemann DA, Strohmeier D, Brodbeck C, Parkkonen L, Hämäläinen MS (2014) MNE software for processing MEG and EEG data. NeuroImage 86:446–460.

Guenther FH, Ghosh SS, Tourville JA (2006) Neural modeling and imaging of the cortical interactions underlying syllable production. Brain Lang 96:280–301.

Harutyunyan A, Dabney W, Mesnard T, Gheshlaghi Azar M, Piot B, Heess N, van Hasselt HP, Wayne G, Singh S, Precup D, Munos R (2019) Hindsight Credit Assignment. In: Advances in Neural Information Processing Systems. Curran Associates, Inc. Available at: https://proceedings.neurips.cc/paper_files/paper/2019/hash/195f15384c2a79cedf293e4a847ce85c-Abstract.html [Accessed July 17, 2024].

Haufe S, Meinecke F, Görgen K, Dähne S, Haynes J-D, Blankertz B, Bießmann F (2014) On the interpretation of weight vectors of linear models in multivariate neuroimaging. NeuroImage 87:96–110.

Howell P, Archer A (1984) Susceptibility to the effects of delayed auditory feedback. Percept Psychophys 36:296–302.

Huth AG, de Heer WA, Griffiths TL, Theunissen FE, Gallant JL (2016) Natural speech reveals the semantic maps that tile human cerebral cortex. Nature 532:453–458.

Issa MF, Khan I, Ruzzoli M, Molinaro N, Lizarazu M (2024) On the speech envelope in the cortical tracking of speech. NeuroImage 297:120675.

Joris PX, Smith PH, Yin TCT (1998) Coincidence Detection in the Auditory System: 50 Years after Jeffress. Neuron 21:1235–1238.

Kakade SM (2001) A Natural Policy Gradient. In: Advances in Neural Information Processing Systems. MIT Press. Available at: https://proceedings.neurips.cc/paper_files/paper/2001/hash/4b86abe48d358ecf194c56c69108433e-Abstract.html [Accessed June 6, 2024].

Kayser J, Tenke CE (2015) On the benefits of using surface Laplacian (current source density) methodology in electrophysiology. Int J Psychophysiol 97:171–173.

Kim KS, Wang H, Max L (2020) It’s About Time: Minimizing Hardware and Software Latencies in Speech Research With Real-Time Auditory Feedback. J Speech Lang Hear Res 63:2522–2534.

Legaspi R, Toyoizumi T (2019) A Bayesian psychophysics model of sense of agency. Nat Commun 10:4250.

Maris E, Oostenveld R (2007) Nonparametric statistical testing of EEG-and MEG-data. J Neurosci Methods 164:177–190.

Matell MS, Meck WH (2004) Cortico-striatal circuits and interval timing: coincidence detection of oscillatory processes. Cogn Brain Res 21:139–170.

Mindlin GB (2013) The physics of birdsong production. Contemp Phys 54:91–96.

Moll FW, Kranz D, Corredera Asensio A, Elmaleh M, Ackert-Smith LA, Long MA (2023) Thalamus drives vocal onsets in the zebra finch courtship song. Nature 616:132–136.

Mulliken GH, Musallam S, Andersen RA (2008) Forward estimation of movement state in posterior parietal cortex. Proc Natl Acad Sci 105:8170–8177.

Nichols TE, Holmes AP (2002) Nonparametric permutation tests for functional neuroimaging: A primer with examples. Hum Brain Mapp 15:1–25.

Oganian Y, Chang EF (2019) A speech envelope landmark for syllable encoding in human superior temporal gyrus. Sci Adv 5:eaay6279.

Oldfield RC (2013) Edinburgh Handedness Inventory. Available at: 10.1037/t23111-000 [Accessed July 16, 2024].

Patil A, Huard D, Fonnesbeck CJ (2010) PyMC: Bayesian Stochastic Modelling in Python. J Stat Softw 35:1–81.

Pedregosa F, Varoquaux G, Gramfort A, Michel V, Thirion B, Grisel O, Blondel M, Prettenhofer P, Weiss R, Dubourg V, Vanderplas J, Passos A, Cournapeau D, Brucher M, Perrot M, Duchesnay É (2011) Scikit-learn: Machine Learning in Python. J Mach Learn Res 12:2825–2830.

Perl YS, Arneodo EM, Amador A, Goller F, Mindlin GB (2011) Reconstruction of physiological instructions from Zebra finch song. Phys Rev E 84:051909.

Pernet CR, Appelhoff S, Gorgolewski KJ, Flandin G, Phillips C, Delorme A, Oostenveld R (2019) EEG-BIDS, an extension to the brain imaging data structure for electroencephalography. Sci Data 6:103.

Rothauser EH (1969) IEEE recommended practice for speech quality measurements. IEEE Trans Audio Electroacoustics 17:225–246.

Schneider DM, Mooney R (2018) How Movement Modulates Hearing. Annu Rev Neurosci 41:553–572.

Schneider DM, Sundararajan J, Mooney R (2018) A cortical filter that learns to suppress the acoustic consequences of movement. Nature 561:391–395.

Sharpee TO, Nagel KI, Doupe AJ (2011) Two-dimensional adaptation in the auditory forebrain. J Neurophysiol 106:1841–1861.

Simonyan K, Horwitz B (2011) Laryngeal Motor Cortex and Control of Speech in Humans. Neurosci Rev J Bringing Neurobiol Neurol Psychiatry 17:197–208.

Sitt JD, Amador A, Goller F, Mindlin GB (2008) Dynamical origin of spectrally rich vocalizations in birdsong. Phys Rev E 78:011905.

Smith SM, Nichols TE (2009) Threshold-free cluster enhancement: Addressing problems of smoothing, threshold dependence and localisation in cluster inference. NeuroImage 44:83–98.

Sutton RS, Barto AG (2018) Reinforcement learning: An introduction. MIT press.

Theunissen FE, David SV, Singh NC, Hsu A, Vinje WE, Gallant JL (2001) Estimating spatio-temporal receptive fields of auditory and visual neurons from their responses to natural stimuli. Netw Comput Neural Syst 12:289–316.

Titze IR (1988) The physics of small-amplitude oscillation of the vocal folds. J Acoust Soc Am 83:1536–1552.

Titze IR (1998) Measurements for voice production: research and clinical applications. J Acoust Soc Am 104:1148.

Tourville JA, Guenther FH (2011) The DIVA model: A neural theory of speech acquisition and production. Lang Cogn Process 26:952–981.

Vehtari A, Gelman A, Gabry J (2017) Practical Bayesian model evaluation using leave-one-out cross-validation and WAIC. Stat Comput 27:1413–1432.

Veillette JP, Nusbaum HC (2025) Bayesian p-curve mixture models as a tool to dissociate effect size and effect prevalence. Commun Psychol 3:9.

Wolpert DM, Diedrichsen J, Flanagan JR (2011) Principles of sensorimotor learning. Nat Rev Neurosci 12:739–751.

Wolpert DM, Ghahramani Z (2000) Computational principles of movement neuroscience. Nat Neurosci 3:1212–1217.

Yamamoto K, Kawabata H (2014) Adaptation to delayed auditory feedback induces the temporal recalibration effect in both speech perception and production. Exp Brain Res 232:3707–3718.

